# Whole-organ surface mapping using multiview projection reconstruction

**DOI:** 10.64898/2026.08.26.747115

**Authors:** Elizabeth Brewer, Milad Almasian, Alireza Saberigarakani, Dongyu Liu, Adam Azizi, Sarah A. Ware, Kunal Karambelkar, Nimit Shah, Meghana Vadlamudu, Girgis Obaid, Dan Tong, Yichen Ding

**Affiliations:** The University of Texas at Dallas, Department of Bioengineering, Richardson, TX, USA; The University of Texas Southwestern Medical Center, Department of Internal Medicine, Dallas, TX, USA; The University of Texas Southwestern Medical Center, Harry Moss Heart Center, Dallas, TX, USA; The University of Texas Southwestern Medical Center, Hamon Center for Regenerative Science and Medicine,Dallas, TX, USA; The University of Texas Southwestern Medical Center, Department of Molecular Biology, Dallas, TX, USA; The University of Texas at Dallas, Center for Imaging and Surgical Innovation, Dallas, TX, USA

**Keywords:** light-sheet, microscopy, cardiac, lymphatic, spheroid, multiview, registration, surface imaging

## Abstract

**Significance:** While light-sheet microscopy is emerging as a robust method for volumetric imaging with improved axial resolution, its capability regarding two-dimensional, surface-level mapping is often hindered by limitations in data redundancy and reconstruction efficiency stemming from volumetric registration methods. We demonstrate that a multiview imaging approach in an axially-swept, dithered light-sheet microscope paired with computational image reconstruction of view projections is able to address these trade-offs to enable large-scale mapping of surface structural features, leveraging the advantages of multiview light-sheet in scalable field of view, working distance, and near isotropic resolution across the entire imaging depth.

**Aim:** To aid in the acquisition and analysis of two-dimensional surface structures, we present a tailored surface mapping workflow and a Fiji plugin for computational reconstruction, promoting robust and comprehensive visualization of surface features of uncleared volumetric samples.

**Approach:** Our strategy, termed projection reconstruction for imaging surface morphology (PRISM), integrates axially-swept dithered light-sheet microscopy and post-processing software for multiview imaging. The imaging hardware enables near-isotropic resolution across its entire field of view, while the software implementation leverages rigid and affine transformations to align two-dimensional projections of multiview samples. It is designed to work with the BigStitcher pipeline, leveraging its robust algorithm to provide support for two-dimensional image alignment and stitching.

**Results:** We demonstrate the capability of PRISM in studies of lymphatic network mapping in the epicardial layer of intact mouse hearts, as well as surface profiles of FaDu spheroids labeled with antibody-nanodiamond conjugates. This method allows us to quantify cardiac lymphatic branch numbers, diameters, and lengths of a Prox1-tdTomato mouse cardiac model, as well as cluster number and diameters of epidermal growth factor receptor within a FaDu spheroid labeled with a nanodiamond-antibody conjugate, with a significant reduction of post-processing data size.

**Conclusions:** PRISM leverages multiview image projections to promote studies of cardiac lymphatics in mouse models and surface receptor distributions within spheroid models, enabling efficient surface mapping of large, intact, and uncleared biological samples across a variety of scales.

## 1 Introduction

Mapping two-dimensional (2D) surface structures across entire intact biological samples presents a unique challenge in optics due to limitations in field-of-view, working distance constraints, lateral and axial resolution inequality, and tradeoffs between resolution and depth of field, which limits the acquisition of high-resolution data across depth. These issues are particularly pronounced in large specimens with dense and heterogeneous structures, where restricted depth of field and photodamage associated with confocal microscopy and other point-scanning approaches hinder surface imaging viability^1,2^. While tissue clearing methods are known to enhance optical transparency and enable deeper interrogation of biological samples by reducing scattering and homogenizing refractive indices^2,3^, significant challenges remain, as current clearing approaches often involve trade-offs between optical transparency, fluorescent protein compatibility, and sample size preservation^4^. As a result, resolving fine surface features throughout the entire organ while maintaining a large field-of-view remains challenging.

Light-sheet microscopy has emerged as a promising strategy for large-scale structural imaging by reducing phototoxicity, providing true optical sectioning, and relaxing the tradeoff between resolution and depth of field^1,2,5^. Leveraging and building upon these advantages, axially-swept light-sheet methods^2,6,7^ directly address disparities between a system’s lateral and axial resolution by exclusively capturing signal at the focal region of the light sheet, where both lateral and axial resolutions are optimal and, under certain parameters, equivalent. In particular, axially swept dithered light-sheet (AS-DiLS) microscopy^8^ extends the confocal region while preserving optical sectioning, enabling near-isotropic resolution across large imaging volumes. In AS-DiLS, imaging performance is constrained by light attenuation throughout uncleared samples, limiting both the illumination and detection geometry such that only regions simultaneously close to the illumination plane and incoming detection focus are captured with high quality. This attenuation thus prevents a single view from providing uniform coverage across the entire specimen, requiring the acquisition of data from multiple angular views. For the application of surface-level optical imaging of uncleared tissues, we present the projection reconstruction for imaging surface morphology (PRISM) strategy, which leverages multiview acquisition^9,10^ to improve coverage and reduce orientation-dependent degradation for mapping of the intact surface features before reconstructing the views to create a map of the entire surface of the specimen. By registering and fusing multiview-acquired 2D projections, PRISM enables organ-wide mapping of intact surface features. To support enhanced reconstruction, we have developed PRISM’s computational framework, available as a Fiji-supported plugin, for efficient and accurate formation of multiview surface datasets with decreased data redundancy. This post-processing platform enables comprehensive, high-resolution visualization of lymphatic networks in intact mouse hearts, particularly at the epicardial layer, as well as nanodiamond-antibody conjugate labeled epidermal growth factor receptor (EGFR) surface profiles in human head and neck squamous cell cancer (FaDu) spheroids.

## 2 Materials and Methods

### 2.1 PRISM-enabled Imaging

Photon attenuation from refractive index inhomogeneity within biological specimens remains a major limitation in conventional optical imaging, restricting the penetration depth in biological tissues to hundreds of micrometers^2–4,11^. Despite the utilization of multiview light-sheet microscopy in tandem with tissue clearing^9,10^, the enhanced volumetric imaging capability is inefficient for the imaging of surface structures due to redundant or irrelevant data acquisition and time-consuming registration, fusion, and reconstruction. For non-cleared samples, optical coverage is limited by the overlap between illumination and detection, typically confining imaging to a restricted angular coverage of less than 90° of the sample **(Fig. 1A)**. To overcome this limitation, multiview imaging increases angular coverage by acquiring images from multiple orientations by rotating the sample relative to the imaging objectives. Regions that are degraded, attenuated, or obscured in one view can thus be complemented by others to reduce orientation-dependent image degradation through computational fusion **(Fig. 1B)**. As the focus of the PRISM platform is to recover the surface of biological samples, we simplify the framework by condensing the acquired volumetric data into maximum intensity projections (MIPs). To prevent the resulting geometric distortion that would result from this dimensionality reduction, we computationally rotate the image volumes back from their relative shift, aligning with the reference view (e.g. 0°). In this work, we employ a 45-degree rotational increment to provide sufficient overlap between adjacent views and ensure robust multiview reconstruction across the entire sample **(Fig. 1C)**. To ensure optimal alignment and image fusion at high resolution, isotropic resolution is required, as each view contributes lateral and axial information to their respective MIPs after the rotational correction. Using AS-DiLS, we achieve a lateral resolution of 4.59 ± 0.59 µm and 4.64 ± 0.58 µm in x and y respectively, and an axial resolution of 4.45 ± 0.62 µm (mean ± s.d., n = 826 beads), enabling consistent near-isotropic resolution across the reconstructed dataset^8^ **(Fig. 1D)**.

**Fig. 1.**
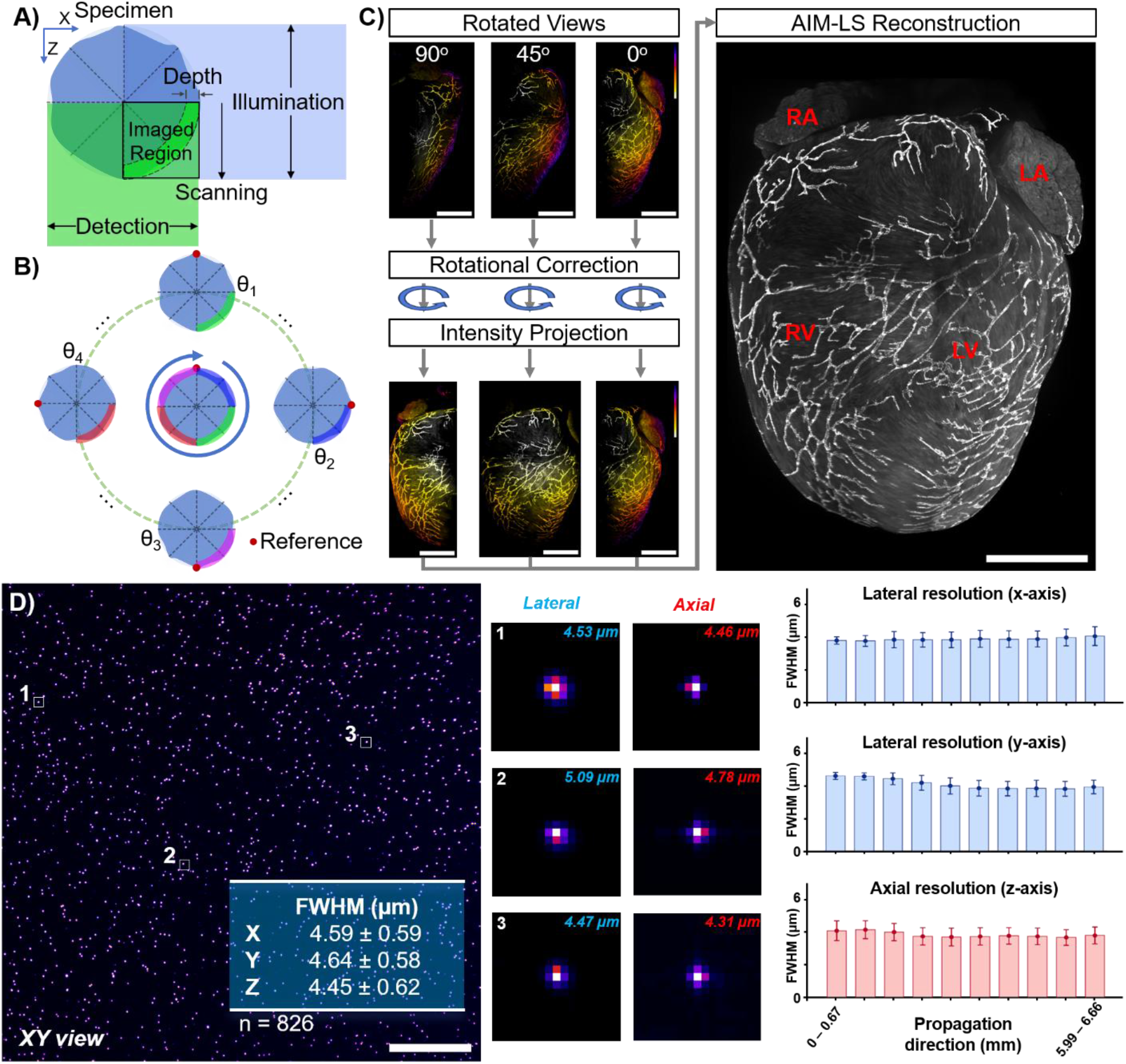
Schematic of the PRISM Strategy: **A)** Diagram demonstrating the illumination/detection penetration depth, which limits the acquisition range of AS-DiLS to less than 90° with respect to a single angular view. **B)** Schematic demonstrating the recoverable areas available to the imaging system given by rotating the sample by some *θ*_*n*_ before imaging. **C)** Overview of the PRISM pre-processing pipeline, covering the acquisition of rotated views, rotational correction, maximum intensity projection, and computational reconstruction with subsequent fusion. Images on left are depicted as depth-encoded projections. Scale bar: 2 mm, Color bar: 0 mm (white) to 4 mm (purple), LA: left atrium, RA: right atrium, LV: left ventricle, RV: right ventricle. **D)** Resolution of AS-DiLS evaluated by fluorescent beads, demonstrating consistent resolving power across the entire field-of-view of the system. Scale bar: 350 µm.

### 2.2 Multiview Reconstruction Plugin

To efficiently reconstruct the surface of a sample from multiple angular views, we have established a computational pipeline to automatically register and fuse projections of the resulting views, creating a cohesive, contiguous surface map of biological samples through customization of the BigStitcher framework^12^ for the purpose of computationally efficient 2D surface mapping. We further developed a user-friendly Fiji plugin to promote the broad application of this pipeline. The pipeline leverages a multiview dataset that allows a portion of overlap (e.g. 10%) between adjacent sample rotations, providing redundant data to identify interest points for registration, that have undergone rotational correction and subsequent 2D projection. While rotational correction accounts for major discrepancies between views, misalignments and minor geometric discrepancies can still occur, limiting the achievable reconstruction quality from standard rigid registration methodologies. Thus, to correct for mismatch between views, scale-invariant feature transformation (SIFT)^13^ is applied to automate the view registration process by computationally optimizing parameters governing rigid and shear translations and scaling coefficients, creating a matrix of matched interest points between respective images. To ensure robusticity during computation, random sample consensus (RANSAC)^14^ is utilized to select the group of inliers identified by SIFT within each set of interest points. For the purpose of having known surface features and geometry, we simulated a spherical 2D shell comprised of long filaments and spherical features across its surface before simulating acquisition and preprocessing prior to PRISM-based fusion. PRISM’s computational workflow begins by loading in 2D MIPs of adjacent views to register **(Fig. 2A)**. One view will be selected to act as the reference upon which the other view will be registered, simplifying the parameter optimization process. The algorithm then performs SIFT-based interest point detection to identify similar structures within the overlapping portions of the views, creating paired points of reference to align **(Fig. 2B)**. This process establishes a set of linear equations **(Fig. 2C)** with respect to the relative coordinates of interest points *A* and the components of the transform matrices *p*_*x*_ and *p*_*y*_, respectively:

**Fig. 2.**
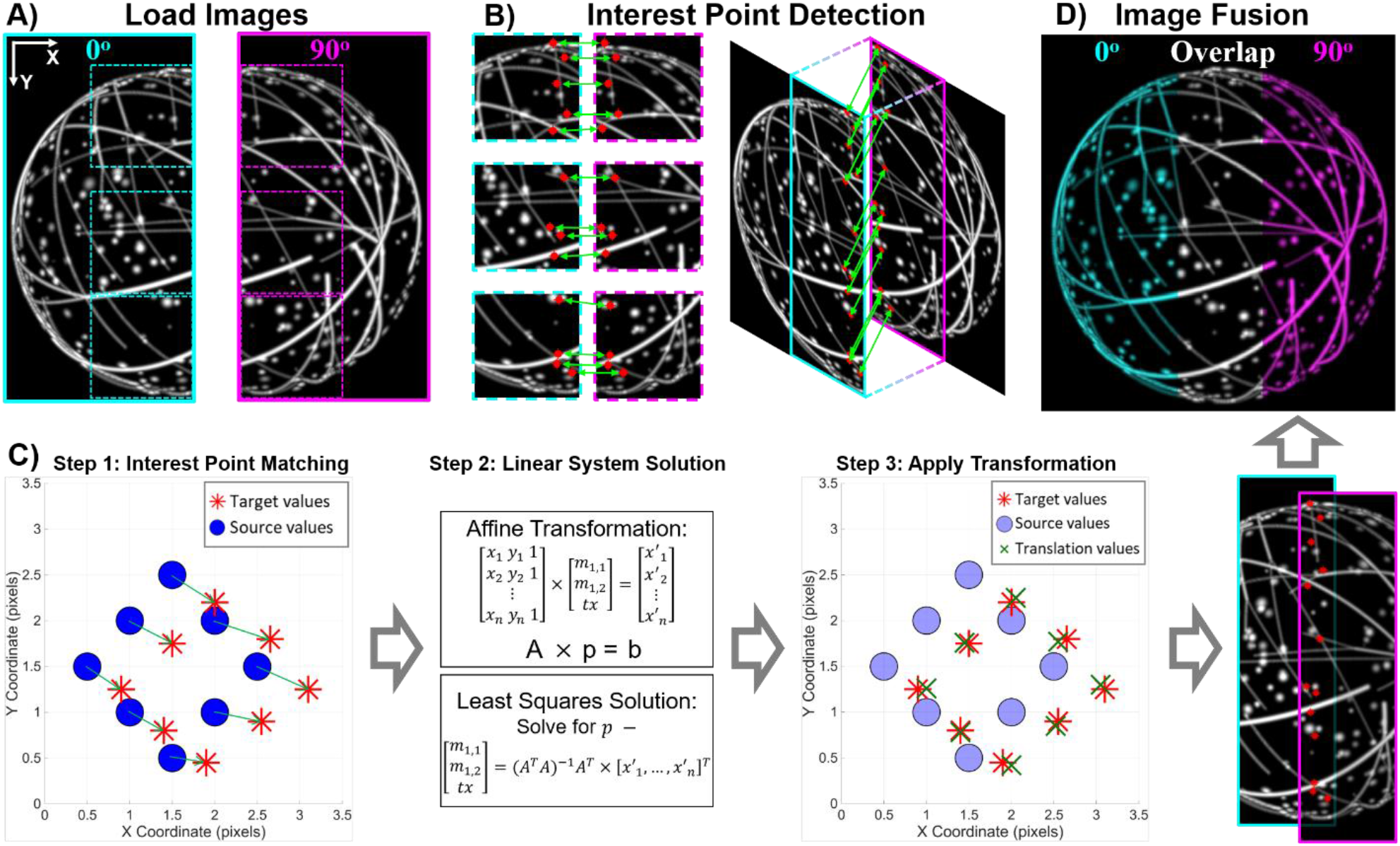
Overview of Multiview Surface Imaging Reconstruction: **A)** The views are loaded as MIPs, where one view acts as the reference (cyan) and the overlapping region between it and the adjacent view (magenta) provides context for registration. **B)** SIFT interest point detection identifies similar structures within the overlapping portions of both views. **C)** The affine transform iteratively adjusts shear (rotation), translation, and scale factors, optimizing the affine transform matrix according to the geometric distance error present between interest points. For brevity, only the case for *x*^′^_1_ is shown. **D)** The reference and registered views are then fused into a single 2D image. The overlapping region is depicted here in white. Additional views can be computed simultaneously or added to the reconstruction by repeating **A-D**.

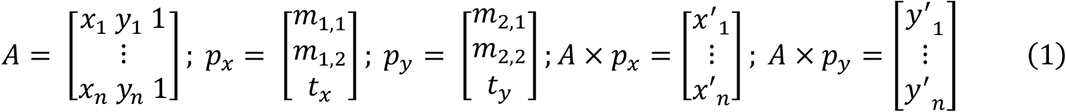

where *A* is the matrix of detected interest points, (*x*_*n*_, *y*_*n*_) and (*x*^′^_*n*_, *y*^′^_*n*_) are the unaligned and reference coordinates, respectively, of the n^th^ registered interest point, *m*_1,1_ and *m*_2,2_ are scaling factors, *m*_1,2_ and *m*_2,1_ are shear (rotational) factors, and *t*_*x*_ and *t*_*y*_ are translational values in x and y, respectively. Least squares optimization then leverages the remaining inlying interest points to minimize root mean squared error and solve for the components of affine transform matrices *p*_*x*_ and *p*_*y*_ as follows:

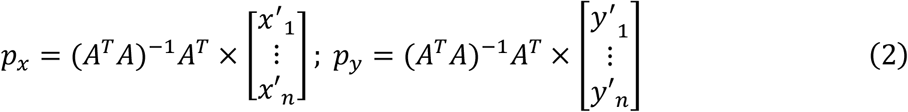

Once the optimal values for *p*_*x*_ and *p*_*y*_ are found, the resulting transformation is applied, and the views are fused into a cohesive image **(Fig. 2D)**.

### 2.3 Sample Preparation and Imaging

The *Prox1-tdTomato* transgenic mouse heart was excised and fixed in 4% paraformaldehyde (PFA; 15714-S, Electron Microscopy Sciences) for 1 h before three 15 min PBS washes. The heart was then mounted in a 2% agarose (16520050, Thermo Fisher Scientific) block, to be imaged with a 2×/0.25NA objective lens (MV PLAPO, Olympus) and zoom body (0.63–6.3×, MVX-ZB10, Olympus) with a 532 nm laser (LRS-0532-PFM-00100-05, Laserglow Technologies). The FaDu human head and neck cancer (ATCC) spheroids were immersed in a suspension consisting of 100 µL of 50 µg/mL nanodiamond-cetuximab-Alexa Fluor 488 *N*-hydroxysuccinimide ester conjugate (nanodiamond: NDNV100nmHiSA2mg, ADAMAS nanotechnology; cetuximab: ERBITUX, Eli Lilly; Alexa Fluor: AF488-NHS, Fisher Scientific). Following this, the spheroids were fixed in 10% formalin for 10 minutes, washed three times with Dulbecco’s phosphate buffer saline (DPBS; without Ca^+2^ and Mg^+2^; Corning), and mounted in a fluorinated ethylene propylene tube with 2% agarose. Afterwards, the tube was immersed in water within the AS-DiLS’ custom chamber before imaging nanodiamond fluorescence with a 12.6×/0.5NA objective lens and zoom body with a 532 nm laser.

## 3 Results

### 3.1 Prox1-tdTomato Transgenic Mouse Lymphatic Vasculature

To highlight the utility of PRISM in biological studies, we imaged the epicardium of an intact heart of a *Prox1-tdTomato* transgenic mouse, targeting its lymphatic vasculature^15^. Cardiac lymphatic vessels play a large role in fluid regulation, immune trafficking, and myocardial remodeling, regulating interstitial fluid clearance and preventing edema^16,17^. They directly influence ventricular compliance and diastolic function, where impaired drainage leads to fluid accumulation and subsequent fibrosis and tissue stiffening, as seen in heart failure with preserved ejection fraction^18^. Despite these critical functions, the global organization of the cardiac lymphatic network remains poorly resolved due to its structural complexity, including region-dependent density, hierarchical branching, and anisotropic alignment along myocardial fibers across epicardial regions. These features are inherently 3D and cannot be accurately captured by serial slicing methodologies, as tissue sectioning disrupts vessel continuity, underestimates connectivity,and fails to resolve volumetric properties such as diameter variation and branching geometry^8^. Thus, accurate characterization requires intact-organ imaging approaches that achieve capillary-scale resolution while preserving whole-heart structural context. Notably, cardiac lymphatic vessels are predominantly distributed within the epicardial layer^15^, well within the available penetration depth of conventional optics. Thus, 2D surface mapping of lymphatic vasculature within the *Prox1-tdTomato* cardiac model can be achieved by leveraging the PRISM post-processing strategy. To this end, eight rotational views of 45° were acquired with our in-house AS-DiLS microscope and fused using the PRISM plugin, with its performance verified by physicians and professionals^15^. The anterior and posterior sides of the heart were isolated to prevent overlapping in the final dataset. Pixel classification using ilastik^19^ was then performed to segment the lymphatic vasculature within the fused image, and filament tracing was used to map the vasculature by node and vessel **(Fig. 3A)**. The anterior side of the heart yielded a branch count of 953, with an average diameter of 18.28 ± 6.59 µm and length of 211.23 ±199.42 µm, and the posterior side yielded 1059 branches of diameter 18.53 ± 7.55 µm and length 188.82 ± 178.40 µm (**Fig. 3B**, data reported as mean ± s.d.). Following these measurements, the diameter of the vessels was then mapped across the anterior side of the mouse heart for visualization of regional variance **(Fig. 3C)**. We thus demonstrate the capabilities of PRISM for the study of cardiac lymphatics across the entire intact epicardium of the *Prox1-tdTomato* mouse heart.

**Fig. 3.**
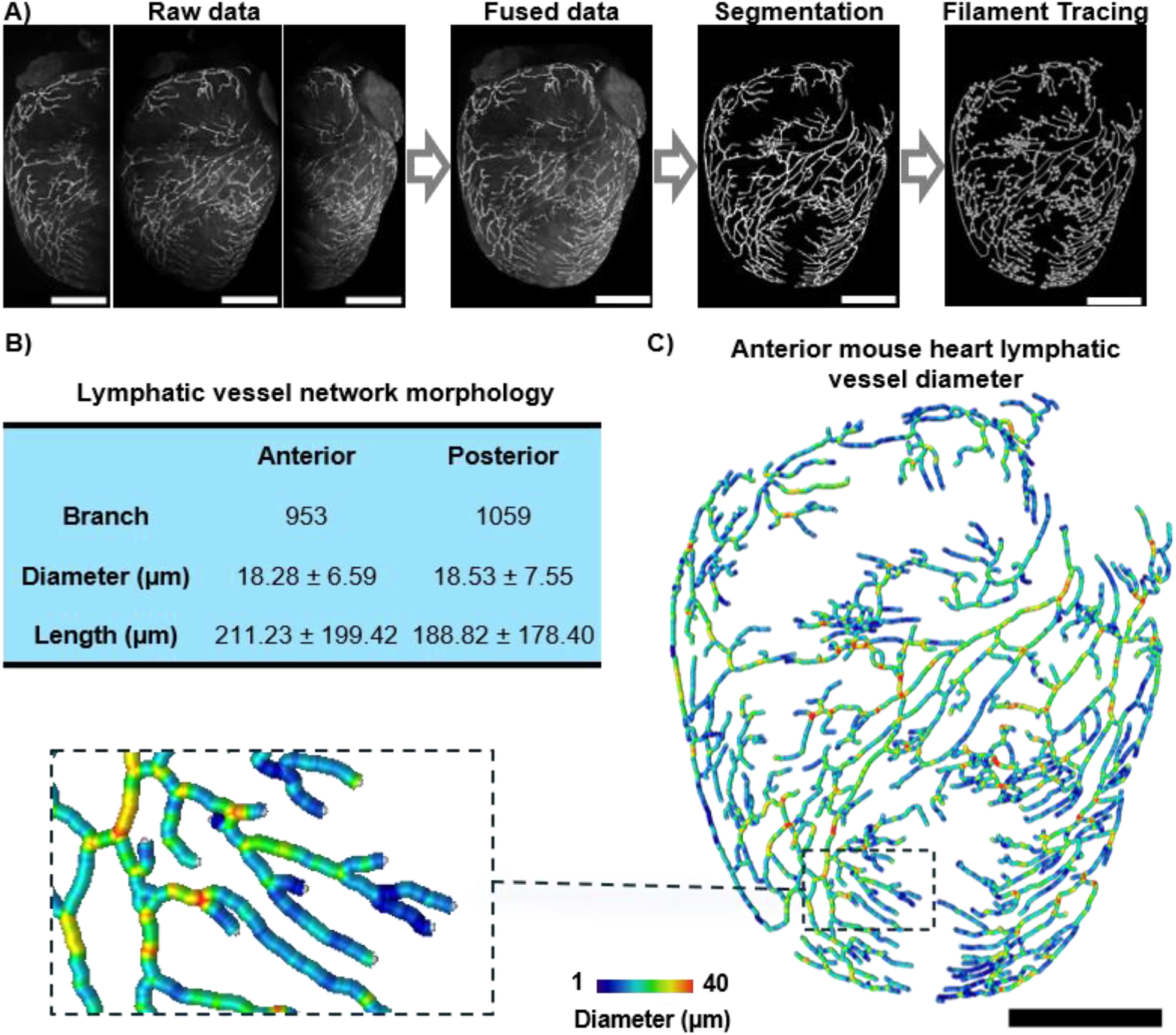
Results of intact cardiac lymphatics investigation using PRISM. **A)** Image processing pipeline for lymphatics interrogation, including data acquisition followed by fusion using PRISM, followed by vessel segmentation and filament tracing for image analysis. Three images are shown for simplicity. **B)** Relevant statistics surrounding lymphatic vessel morphology derived from the entire intact *Prox1-tdTomato* mouse heart model. **C)** Colormap of the anterior lymphatic vasculature, demonstrating size variations across the entire mouse heart. Scale bar: 2 mm.

### 3.2 Nanodiamond-antibody Conjugate Labeled EGFR in FaDu Spheroid

We further demonstrate the capabilities of PRISM by imaging EGFR distributions on the surface of FaDu spheroids labeled with antibody-conjugated fluorescent nanodiamonds, as per our expertise^20^. Nanodiamond probes for fluorescence imaging are known for their low intrinsic cytotoxicity^21^ and excellent photostability^22^, making it a good candidate for multiview cellular imaging, as repeated measurements can increase the risk of photobleaching in organic dyes. To increase biocompatibility, nanodiamonds were treated with a streptavidin coating before conjugation with a cetuximab-AF488-biotin antibody complex to enable EGFR labeling. After immunolabeling, the FaDu spheroids were fixed in 10% formalin for 10 minutes at room temperature. Given the relatively small size of the FaDu spheroid (∼600 µm in diameter) in comparison with our field of view (∼1 mm), whole coverage of one side was able to be achieved in two rotational views spaced 45 degrees apart **(Fig. 4A)**. After fusion through the PRISM plugin **(Fig. 4B)**, these two views were manually segmented into nanodiamond-labelled clusters **(Fig. 4C)** before further processing to extract the diameter of the clusters. These cluster diameters were mapped according to their respective cluster to show the spatial distribution of cluster size and placed into 10 bins, each 1 µm in width, to quantify the distribution of cluster diameters on the surface of the spheroid **(Fig. 4D)**. With an average diameter of 2.35 ± 0.99 µm (mean ± s.d.), the nanodiamond clusters demonstrate aggregation at subcellular scale, demonstrating the capabilities of PRISM to map surface structures across FaDu spheroids by leveraging multiview rotational acquisition.

**Fig. 4.**
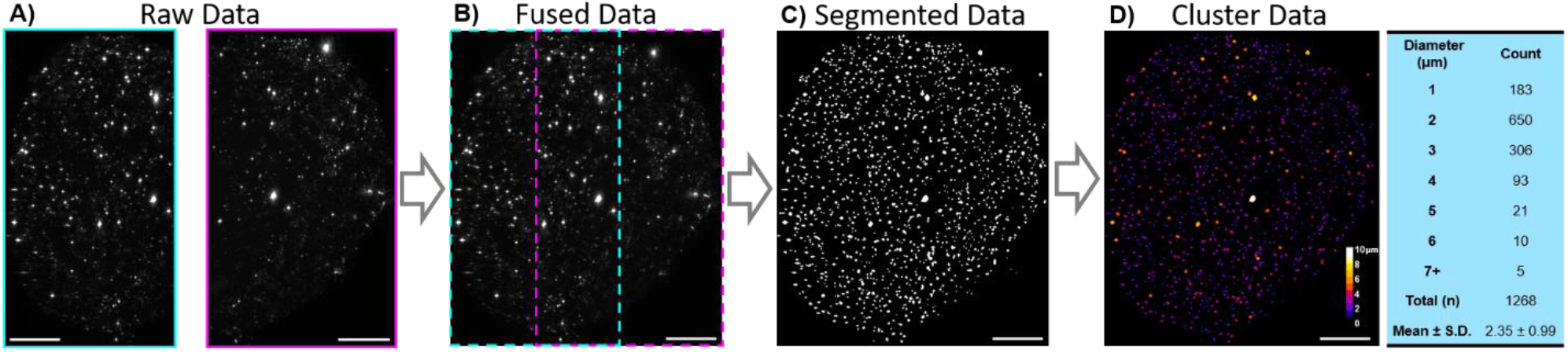
Nanodiamond-antibody labeled FaDu spheroid results. **A)** Raw data from acquisition of 0- (magenta) and 45-degree (cyan) angles. **B)** Fused image following PRISM post-processing. **C)** Segmentation results of nanodiamond clusters on the surface of the spheroid. **D)** Colormap of nanodiamond-labeled spheroid clusters and subsequent table showing cluster diameter distribution. Scale bar: 100 µm.

## 4 Discussion

We present the PRISM strategy and Fiji plugin for 2D surface mapping of intact biological samples, demonstrating its capabilities through epicardial lymphatic imaging of whole *Prox1-tdTomato* transgenic mouse hearts, as well as EGFR labeling in nanodiamond-antibody conjugated FaDu spheroids. By leveraging a rotational image acquisition scheme to cover the whole surface of the sample before a computational reconstruction plugin that leverages RANSAC-refined SIFT for interest point detection and a dual rigid-affine transformation, we achieve full surface feature mapping of intact biological samples. As a generalized framework for axially-swept light sheet microscopy, whole-mount sample manipulation, including a rotational axis, is required to maximize the performance of PRISM. Additionally, sample geometry may limit achievable acquisition. For example, extreme concave surfaces may restrict illumination/detection geometry, resulting in missing data. Finally, while RANSAC-refined SIFT is a powerful, feature-based registration method, its parameters require fine tuning based on the provided dataset, which may require additional optimization. To aid in this, we provide a window during operation of the PRISM plugin that prompts for specific values of relevant parameters. Future work will go towards implementing further support regarding imaging methodology, increased restoration quality through automated parameter optimization, as well as applications towards further studies into cardiac lymphatics and surface proteins in spheroid models.

## Code and Data Availability

The computational image reconstruction framework, implemented in Java as a Fiji-compatible plugin, can be found on Zenodo (10.5281/zenodo.22085688).

## Acknowledgements

This study was supported by National Institutes of Health grant R00HL148493 (Y.D.), National Institutes of Health grant R01HL162635 (Y.D.), National Science Foundation grant 2503230 (Y.D.), AHA grant 25TPA 1472117 (D.T.), and Cecil H. and Ida Green Professorship in Systems Biology Science (Y.D.).

